# Image recovery from unknown network mechanisms for DNA sequencing-based microscopy

**DOI:** 10.1101/2022.09.29.510142

**Authors:** David Fernandez Bonet, Ian T. Hoffecker

## Abstract

Imaging-by-sequencing methods are an emerging alternative to conventional optical micro- or nanoscale imaging. In these methods, molecular networks form through proximity-dependent association between DNA molecules carrying random sequence identifiers. DNA strands record pairwise associations such that network structure may be recovered by sequencing which, in turn, reveals the underlying spatial relationships between molecules comprising the network. Determining the computational reconstruction strategy that makes the best use of the information (in terms of spatial localization accuracy, robustness to noise, and scalability) in these networks is an open problem. We present a graph-based technique for reconstructing a diversity of molecular network classes in 2 and 3 dimensions without prior knowledge of their fundamental generation mechanisms. The model achieves robustness by obtaining an unbiased sampling of local and global network structure using random walks, making use of minimal prior assumptions. Images are recovered from networks in two stages of dimensionality reduction first with this structural discovery step followed by the manifold learning step. By breaking the process into stages, computational complexity could be reduced leading to fast and accurate performance. Our method represents a means by which diverse molecular network generation strategies could be unified with a common reconstruction framework.

---

Imaging-by-sequencing methods (Zador et al., 2012; Glaser et al., 2015; Schaus et al., 2017; Boulgakov et al., 2018; Weinstein et al., 2019; Hoffecker et al., 2019; Gopalkrishnan et al., 2020; Boulgakov et al., 2020; Greenstreet et al., 2022) arose recently as a potential alternative molecular imaging strategy based entirely on DNA sequencing information in contrast to optical or optics-coupled techniques like spatial omics (Ke et al., 2013; Lee et al., 2015; Ståhl et al., 2016; Wang et al., 2018; Karaiskos et al., 2017; Satija et al., 2015; Achim et al., 2015; Halpern et al., 2017), single molecule localization microscopy (Söderberg et al., 2006; Jungmann et al., 2010; Rust et al., 2006; Betzig et al., 2006), or fluorescence imaging more broadly. Individual nanoscale molecular associations in imaging-by-sequencing lead to unique sequence-based records that denote local proximity between molecules. This notion of proximity-dependent association is extended to form large interconnected networks that encompass a global space (Fig.1 A). We can formally represent this spatial information as a graph, where strand origins are nodes and strand-to-strand associations are edges. An open computational problem is that of optimal image recovery, i.e. the task of how to quickly and accurately reorganize a collection of local pairwise associations obtained from sequencing into a spatial representation reflecting the underlying nano- or microscale molecular distribution.

**Figure 1.**
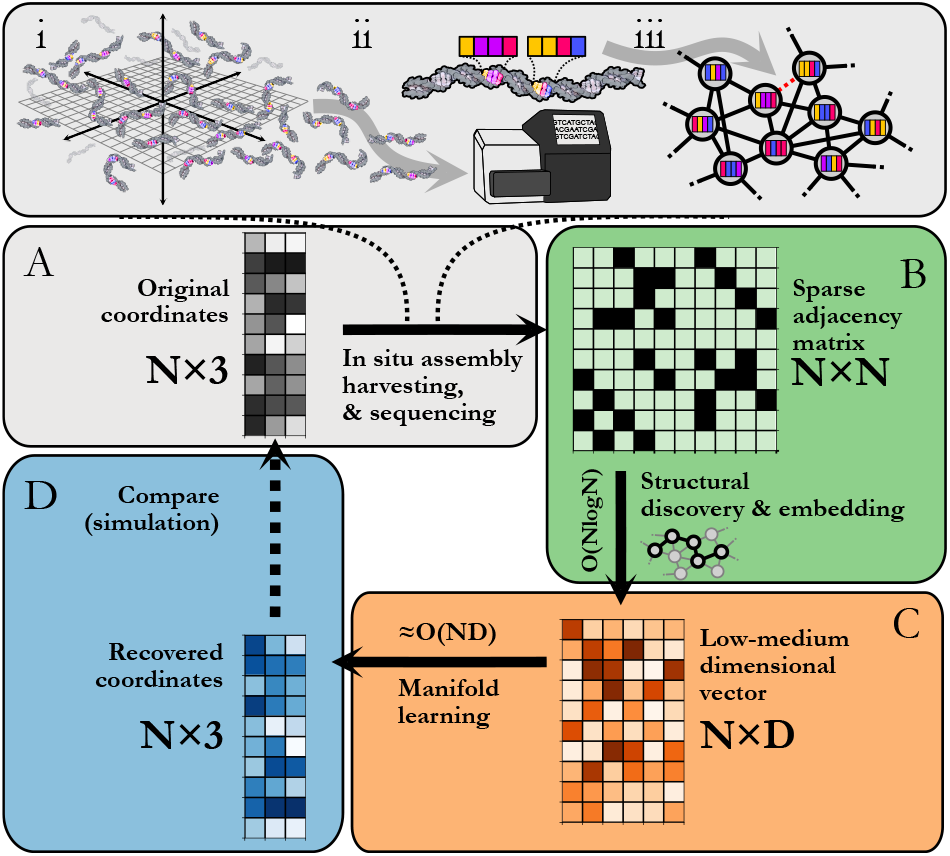
Imaging-by-sequencing by staged graph embedding: A. (i) *in situ* network formation, (ii) harvesting, sequencing, and cataloging of network edges, (iii) re-assembly of an abstract network. B. Beginning from an NxN pairwise binary adjacency matrix, an initial structural discovery step uses random walk sampling to compress the data into an NxD matrix. C. The remaining dimensionality reduction is carried out with manifold learning. D. The recovered Nx3 or Nx2 set of coordinates is an approximation of the original (unknown) set of relative molecular positions.

Imaging-by-sequencing has seen a variety of network generating rules giving rise to different systematic patterns of sub-graphs or network structural motifs. For example: networks where nodes are connected to most other nodes (Weinstein et al., 2019), locally connected Voronoi meshes (Hoffecker et al., 2019), locally connected neighbor networks (Boulgakov et al., 2018; Schaus et al., 2017; Gopalkrishnan et al., 2020; Boulgakov et al., 2020), “GPS” networks (Greenstreet et al., 2022), or boundary-sharing cell networks (Zador et al., 2012; Glaser et al., 2015). Strategies moreover fall into distance-weighted(Weinstein et al., 2019; Gopalkrishnan et al., 2020; Greenstreet et al., 2022) or binary unweighted categories(Schaus et al., 2017; Hoffecker et al., 2019; Boulgakov et al., 2018). These design differences at the fundamental structure level seem to indicate that there cannot be a one-size-fits-all reconstruction strategy. Moreover, a still greater challenge arises when attempting reconstruction while lacking complete knowledge of the microscopic processes driving network formation which are likely more physically complex than their design. In this study, we show how to achieve robust spatial reconstruction from a class of arbitrary network motifs under conditions of minimal prior knowledge of the underlying generating rules.

Following *in situ* self assembly, harvesting, and sequencing of a spatial DNA network (Fig.1 A), our processing pipeline begins from a pairwise adjacency matrix of the form

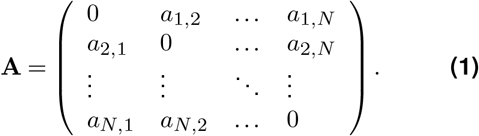

such that

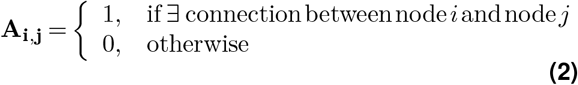

We then reduce the dimensionality of the initial N×N matrix in two stages. First, we perform a structure discovery step based on graph representation learning (Hamilton, 2020), which performs random walks to sample the local and global structural characteristics in the neighborhood of each node in the graph (Node2Vec (Grover and Leskovec, 2016)), a step we shall refer to as structural embedding (Fig.1 B). The output yields an intermediate multi-dimensional vector of the form

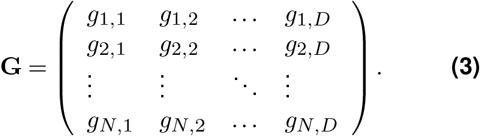

such that typically *D* ≪ *N*. Second, the intermediate vector is fed into a subsequent dimensionality reduction stage (Fig.1 C) that uses manifold learning to embed N points into either 2 or 3 dimensions, i.e. spatial coordinate vectors of the form

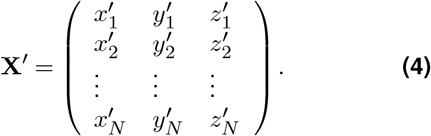

These reconstructed points (Fig.1 D) approximate the original image within some accuracy. All reconstructions are obtained using the hyperparameters in Table 2, which are found to be a compromise between reconstruction accuracy and low computational complexity. For a grid search involving such parameters, refer to Fig.S1.

We use a staged process to reduce computational complexity. Compressing the adjacency matrix to the form in Eq.3 through structural embedding is beneficial as the computational complexity for manifold learning becomes effectively linear ≈ *O*(*N*) since *D* ≪ *N* as will generally be the case for large networks. Furthermore, structural embedding has an upper bound time complexity *O*(*N*log*N*), a major improvement compared to directly applying manifold learning with complexity *O*(*N*^2^), where *D* = *N*.

Redistributing tasks to reduce overall complexity is a strategy that is also employed in an aesthetic graph drawing method (Harel and Koren, 2002), whereby shortest-path distances from key nodes are computed for structural embedding, and subsequent reduction is carried out with principal component analysis (Wold et al., 1987) (PCA). Our method differs in that random walk structural embedding is less computationally demanding than determining shortest-path-distances. Furthermore, we desire strict preservation of all geometric relationships, and thus used UMAP(McInnes et al., 2018) over PCA for its superior preservation of local geometry.

Using structural discovery followed by manifold learning, we reconstructed both 2 and 3-dimensional simulated point distributions (Fig.2 A). Initial molecule positions were randomly distributed over the space of a square or cube of characteristic length *L* =1 with no prior knowledge of molecule identity relative to position so as to model the dispersion of DNA in an imaging-by-sequencing experiment.

**Figure 2.**
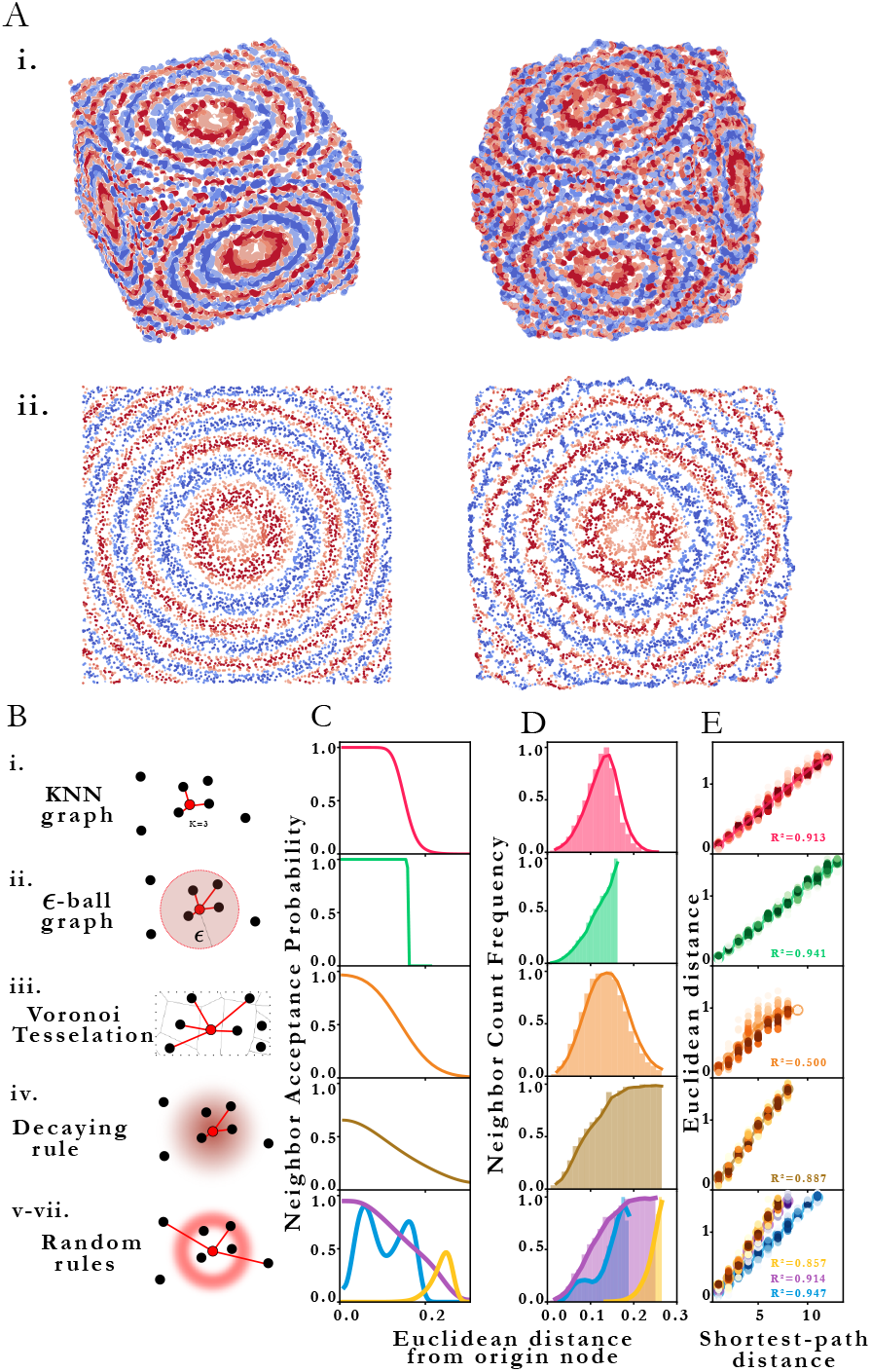
2 stage reconstruction by structural discovery and manifold learning. A. Original (left) set of points and resulting reconstructed (right) points in i. 3 dimensions and ii. 2 dimensions. B. Types of proximity graphs (i-vii) and illustrated corresponding edge drawing rules, featuring probability clouds for stochastic rules. C. Probability that nodes *i* and *j* are neighbors (share an edge) given that their Euclidean distance is *d_ij_* for each type of proximity graph (i-vii) respectively. D. Neighbor count frequency with distance. E. Correlation between Euclidean distance and Shortest-path distance for each network type for each type of proximity graph (i-vii) respectively.

To explore the method’s robustness to variation in network motif, we chose multiple different rule sets for determining how randomly dispersed points assemble into an interconnected network. Each rule can be seen as representing a different model of the physical association process (Fig.2 B), i.e. different mappings of proximity to edges. The different “proximity graph” (Zemel and Carreira-Perpiñán, 2004) definitions are summarized in Table 1. We explored 3 deterministic (KNN-graph, *ε*-ball graph, Voronoi tessellation) and 4 stochastic proximity graphs (based on arbitrary probabilistic rules). For completeness, we also examined a KNN distance-weighted graph in contrast to the binary graphs represented by Eq.2, whereby edges are weighted by some function of separation distance (in this case the inverse distance).

**Table 1.** Proximity graph rules

| Proximity graph | Rule: connect origin node to candidate... |
| --- | --- |
| i. KNN graph | if among the $k$ closest neighbors |
| ii. ε-ball graph | if within distance $\epsilon$ to origin node |
| iii. Voronoi tessellation | if Voronoi cell shares border with origin cell |
| iv. Decaying rule | according a distance-decaying probability |
| v-vii. Random rules | according to arbitrary probability distribution |

**Table 2.** Image reconstruction hyperparameters. Default value and definition

| Parameter | Default value | Definition |
| --- | --- | --- |
| neighbors | 15 | [UMAP] average number of neighbors per node |
| embedded dimension | 32 | [Node2Vec] each node is mapped into a vector of that many dimensions |
| 1/q | 1 | [Node2Vec] the higher the more Breadth-First-Search alike random walks |
| 1/p | 1 | [Node2Vec] the higher the more Depth-First-Search alike random walks |
| random walk length | 10 | [Node2Vec] number of visited nodes in each random walk |

Network generation rules exhibit characteristic neighbor acceptance probability distributions as a function of distance between neighbor and origin node (Fig.2 C). For an arbitrary set of randomly distributed points, different rules produce distinct neighbor frequency distributions, i.e, (normalized) number of neighbors encountered as a function of distance from a given node (Fig.2 D). We observed that all network rules gave rise to monotonic relationships between the average Euclidean distance and graph shortest-path distance (Fig.2 E). This observation suggests a basis for geometry preservation between Euclidean and graph space - i.e. that there exists a unique expected Euclidean distance corresponding to each shortest-path-distance in a given reconstructed network. A geometric relationship between a set of points represented as a set of shortest-path-distances in graph space may thus be expected to have a corresponding (though probabilistic) relative geometric relationship in Euclidean space due to this mapping.

Ground truth access via simulation enables us to compare original and reconstructed points to assess accuracy. We quantify accuracy according to three standards: a local quality metric, a global quality metric and a mean distortion quality metric. The local quality metric (KNN, Supplementary Section E) examines the difference between original and reconstructed neighborhoods of every point, and is an indicator of the fine structure. Conversely, the global quality metric (CPD, Supplementary Section F) examines the pairwise distance Pearson correlation between original and reconstructed points and is an indicator of the coarse structure. Similar assessments of the local and global structures are common practice when comparing high and low-dimensional data (Kobak and Berens, 2019). Lastly, the mean distortion is obtained by performing an affine transformation (Supplmentary Section G) on the reconstructed points. Distortion simply measures the displacement between original and reconstructed points: the less displacement there is, the better the reconstruction. The mean distortion is obtained by averaging the distortion of all points.

Fig.3 A shows a visualization of distortion following reconstruction of 10000 points for 2D and 3D cases. While central points show below-average distortions, border points seem to exhibit higher distortions. We speculate that this may be attributed to differences in characteristic topology near the boundaries which diverge from the patterns exhibited in the core of the point cloud.

**Figure 3.**
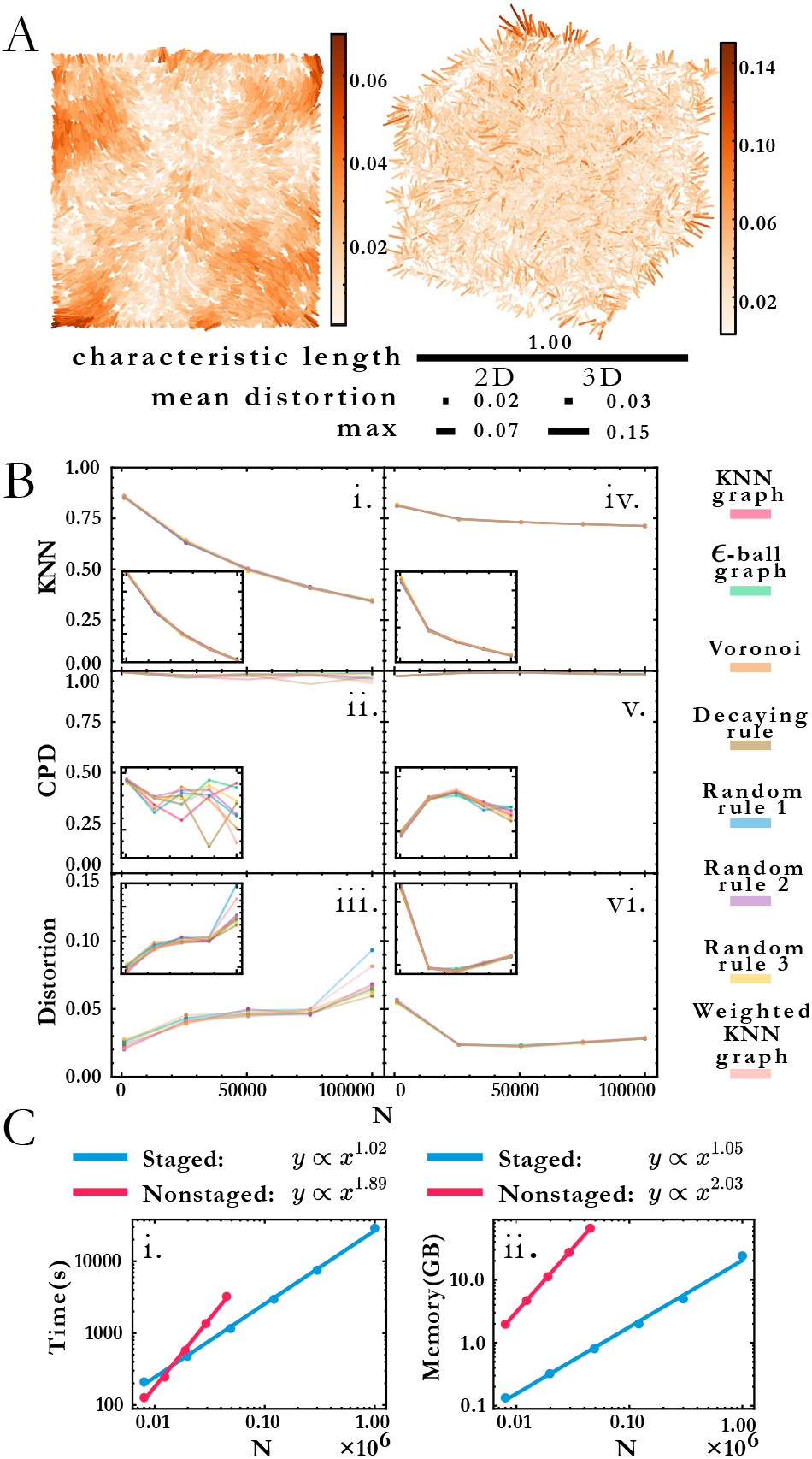
A. Distortion map for the 2D and 3D cases on left and right respectively. Line segments connect original and reconstructed points and are colored according to the displacement (distortion) between them. Relative scale bars for the mean distortion, the maximum distortion and the characteristic length are shown underneath. B. Quality metric dependence with proximity graph type and system size for 2D and 3D respectively: i. and iv. Local quality metric, ii. and v. Global quality metric, iii. and vi. Mean distortion. C. Computational complexity for the staged approach (blue) and the not staged approach (red) for time complexity (left) and memory complexity (right).

Reconstruction accuracy dependence is measured via three parameters: dimension, system size and proximity graph type (Fig.3 B). Interestingly, accuracy in all three categories does not appear to vary much by proximity graph type (weighted or unweighted) as shown in Table 3. Even random proximity rules (e.g. an arbitrary bimodal neighbor acceptance probability) exhibit stable quality trends in line with the other graph types. We observed that accuracy worsens as system size increases, with trends in 3D less severe than 2D. Local reconstruction quality according to the KNN metric for the 2D case (Fig.3 B i.) was robust to proximity graph type, with a maximum variation of 1.5% but diminished with system size. For instance, reconstructed and original points shared less than 40% overlap in neighbors for *N* = 10^5^. Conversely, the 3D case (Fig.3 B ii.) exhibited a gradual decrease with the system size, suggesting that 3D reconstruction tolerates more points. It is also more robust, with a maximum variation of 0.7% between proximity graph types. The global quality metric (Fig.3 B iii-iv.) showed that pairwise distances between original and reconstructed points were linearly correlated (Pearson coefficient is markedly close to 1), indicating that relative distances are preserved during reconstruction. Global quality in the 2D case exhibits the largest variation to proximity graph type, with a maximum of ≈ 6%, whereas the maximum variation in 3D was an order of magnitude lower at 0.6%. Distortion also does not vary much with proximity graph type (Fig.3 B v-vi.). However, in agreement with the other metrics, distortion worsens with increasing points. An exception to this tendency was when system size was small enough in the 3D case (*N* = 1000), with distortion improving within this increment (*N* > 20000). Overall we note that the pipeline works without user supplied knowledge of or adaptation to these structural differences, as this is managed automatically in the structural discovery stage. This would be a major advantage in an experimental setup with imperfect knowledge of the molecular processes leading to proximity associations. The reconstruction accuracy size dependence might be mitigated by adjusting hyperparameters of the structural discovery step, for example greater random walk lengths for larger systems.

**Table 3.** Maximum and mean quality metric variation over the different proximity graphs, for the 2D and 3D case.

|  | Mean variation - 2D | Max variation - 2D | Mean variation - 3D | Max variation - 3D |
| --- | --- | --- | --- | --- |
| KNN | 0.011 | 0.015 | 0.002 | 0.007 |
| CPD | 0.032 | 0.058 | 0.003 | 0.006 |
| Distortion | 0.011 | 0.033 | 0.001 | 0.002 |

We obtained accurate reconstructions from both weighted and binary unweighted designs (Fig.3 B), which is noteworthy as the unweighted designs store less information than their distance-weighted counterparts. This would seem to support the validity of setups that only record whether an interaction happened or not (binary design) versus setups that record a measure of the distance between points (weighted design).

Staged embedding significantly improves computational complexity (Fig.3 C). We compare our approach to direct application of manifold learning to the shortest-path distance matrix (obtained by running a Breadth-First-Search algorithm at the graph in Eq.1). While this nonstaged approach can also reconstruct 2 and 3-dimensional images, its computational complexity becomes prohibitive for a large number of points, both time-wise and memory-wise (Fig.3 C i. and ii.). However, the staged approach tackles the computational complexity issue by effectively compressing the adjacency matrix using the random walk-based structural discovery step (Eq.3). Subsequent manifold learning task becomes less resource-consuming, dealing with a *D*-dimensional vector instead of an *N*-dimensional vector (where *D* ≪ *N*). Whereas the nonstaged approach exhibits near-quadratic empirical scaling *O*(*N*^2^) in both time and memory, the staged approach has near-linear complexity. This should enable large, fast reconstructions. Reconstructing a *N* = 10^6^ image with the nonstaged approach would take years, and reconstructing the same image using the staged approach took eight hours.

Realizing the promise of imaging-by-sequencing will require robust, scalable reconstruction strategies. The method presented here addresses robustness to uncertainty regarding graph generating mechanisms, however the field will also need tools for dealing with systematic variations in network motifs as these might arise in biological imaging scenarios, e.g. anomalously high or low density regions. The problem of scalability will also need to be continuously addressed, as falling sequencing prices enabling greater experiment throughput will push the demand for computational efficiency. Finally, while in this work we made use of quality metrics that compare reconstructed results to those of simulated ground truth data, however it will be important to develop quality metrics that may be used without ground-truth knowledge as will be the case in experimental contexts.

## Author Contributions

DFB implemented the algorithms, characterization, and computational exploration in the study. DFB and ITH conceived the study and wrote the manuscript.

## Acknowledgements

We acknowledge support from the Swedish Research Council (no. 2020-05368), the A. W. Stiftelsen (M19-0547), and the European Research Council ERC (no. 949624 to IH).

## Supplementary Information

### A. Node2Vec

Node2Vec (Grover and Leskovec, 2016) captures graph structural information through random walks, embedding nodes into high-dimensional vectors. It effectively compresses the information of the adjacency matrix into a *N* × *D* matrix (Eq.3). Storing information in such a way improves the overall computational complexity for the subsequent manifold learning and the robustness of our approach. The structural embedding is optimized by maximizing the loglikelihood of observing a node given a particular random walk context. In short, several random walks are performed starting at each node while storing every visited node. This information is used to optimize randomly initialized vectors. Negative sampling is used to reduce the optimization’s computational complexity.

### B. UMAP

UMAP (McInnes et al., 2018) is a manifold learning technique focused on preserving local relations. Because of this, it performs better when dealing with proximity information. It is worth noting that UMAP can directly recover images without requiring a previous structural embedding step, as done in Fig.3 C. A dense version of the adjacency matrix Eq.1 can be obtained by computing the shortest-path distance between every node, where *d_ij_* is the shortest-path distance between node *i* and node *j*. The result is a NxN distance matrix, assigning N-dimensional vectors to each node. Here, the linear relation between Euclidean distance and graph shortest-path distance (Fig.2 E) is exploited to obtain high-dimensional vectors that preserve Euclidean distances.

### C. Hyperparameters

We use the values shown in Table 2 as hyperparameters when using Node2Vec and UMAP. We found them to be a good compromise between reconstruction accuracy and low computational complexity.

We also investigated the influence on the local and global quality metric over deviations from the default parameters (Fig. S1).

### D. The local and global structure

Local structure is preserved when points that are close to each other in the original image are also close in the reconstructed image. In this work, local structure is investigated through the KNN quality metric, which evaluates the fraction of neighbors preserved in the reconstruction. Conversely, global structure is preserved when points that are separated far apart in the original image are equally far in the reconstructed image. In this case, the quantity to be preserved is separation distance, and it can be investigated by computing the correlation between pairwise distances.

### E. KNN

The KNN quality metric can be formally defined as

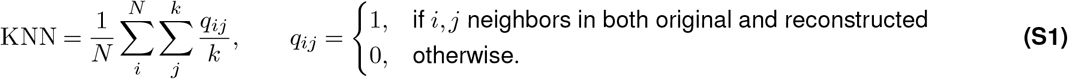

Here, *N* is the total number of elements and *k* is the number of considered neighbors. We used *k* = 15 for quality metrics presented in this manuscript. Effectively, KNN computes the fraction between original and reconstructed neighbors (inner summation) for every node (outer summation) and averages it over the total number of elements.

**Figure S1.**
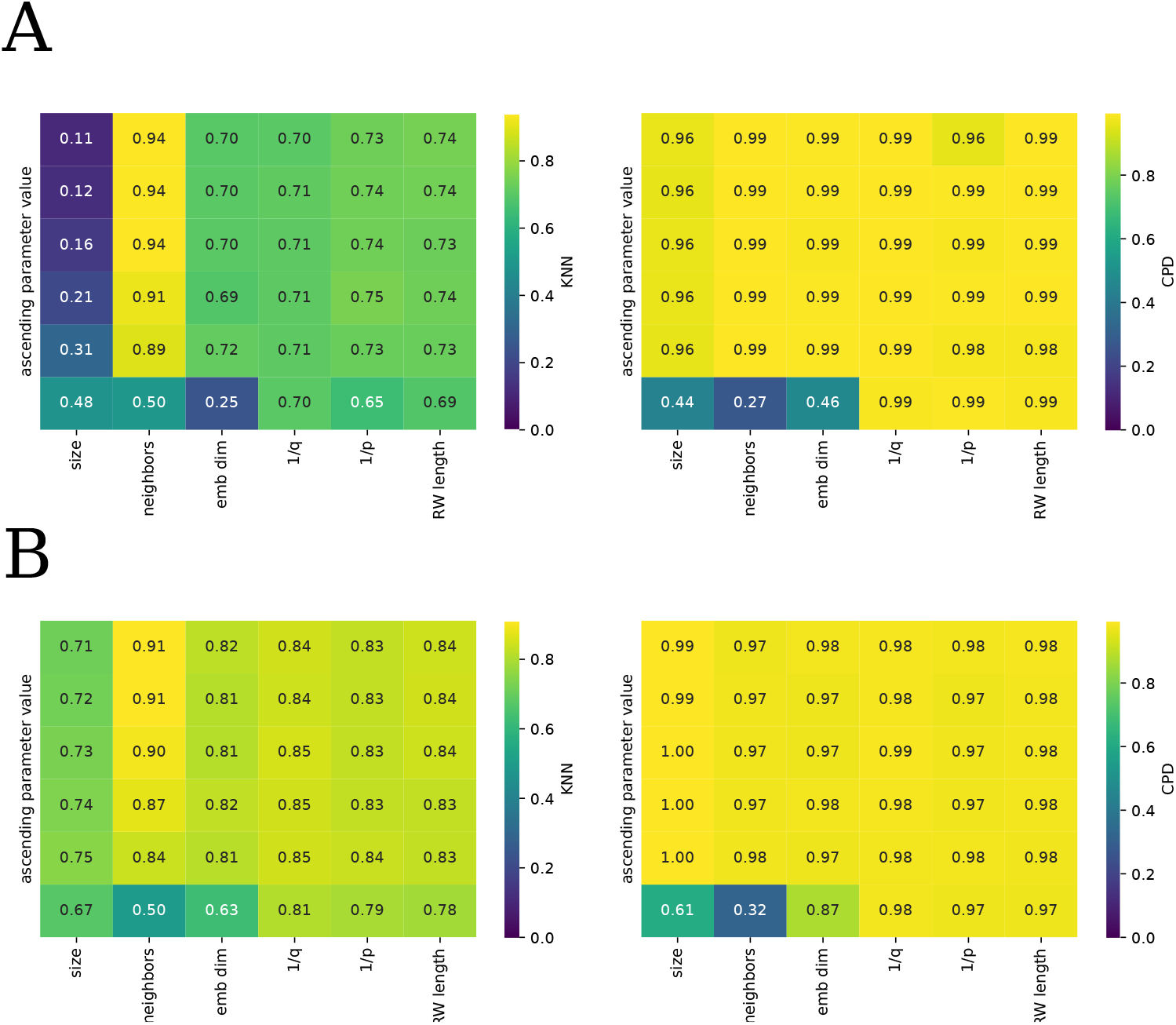
A. Parameter grid search for the 2D case and B. Parameter grid search for the 3D case.

### F. CPD

The CPD quality metric is the Pearson correlation between original and reconstructed pairwise distances. As opposed to KNN, CPD measures concern not only neighboring points but also distant points. The motivation to measure correlation is to evaluate distance preservation. If pairwise distances in the original image are linearly correlated with pairwise distances in the reconstructed image, then the reconstructed image is equivalent to the original image up to an affine transformation. Dealing with scalability often require workarounds: for *N* points, there are 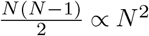 pairwise distances, which can be computationally prohibitive for large *N*. The adopted solution is to randomly sample *N* = 1000 points and compute their 499500 pairwise distances for the original and reconstructed points, to later obtain a representative Pearson correlation.

### G. Distortion

Point cloud alignment is necessary to compute distortion, i.e, the distance between each original and reconstructed point pair. Consequently, we wish to align the original and reconstructed point cloud. To perform such an alignment we apply a affine transformation, i.e, a composition of translation, rotation, scaling and reflection such that distortion is minimized. In particular, we apply the coherent point drift algorithm (Myronenko and Song, 2010). However, if the point clouds are initially far apart, such algorithm becomes computationally demanding. To solve this, we first perform a set of transformations over the reconstructed points involving scaling obtained by computing the correlation between pairwise distances, point-to-point translation and rotation, and the set of coordinate reflections that minimize distortion.

